# Robust cytoplasmic partitioning by solving an intrinsic cytoskeletal instability

**DOI:** 10.1101/2024.03.12.584684

**Authors:** Melissa Rinaldin, Alison Kickuth, Benjamin Dalton, Yitong Xu, Stefano Di Talia, Jan Brugués

## Abstract

Early development across vertebrates and insects critically relies on robustly reorganizing the cytoplasm of fertilized eggs into individualized cells. This intricate process is orchestrated by large microtubule structures that traverse the embryo, partitioning the cytoplasm into physically distinct and stable compartments. Despite the robustness of embryonic development, here we uncover an intrinsic instability in cytoplasmic partitioning driven by the microtubule cytoskeleton. We reveal that embryos circumvent this instability through two distinct mechanisms: either by matching the cell cycle duration to the time needed for the instability to unfold or by limiting microtubule nucleation. These regulatory mechanisms give rise to two possible strategies to fill the cytoplasm, which we experimentally demonstrate in zebrafish and *Drosophila* embryos, respectively. In zebrafish embryos, unstable microtubule waves fill the geometry of the entire embryo from the first division. Conversely, in *Drosophila* embryos, stable microtubule asters resulting from reduced microtubule nucleation gradually fill the cytoplasm throughout multiple divisions. Our results indicate that the temporal control of microtubule dynamics could have driven the evolutionary emergence of species-specific mechanisms for effective cytoplasmic organization. Furthermore, our study unveils a fundamental synergy between physical instabilities and biological clocks, uncovering universal strategies for rapid, robust, and efficient spatial ordering in biological systems.

---

Physical mechanisms play a fundamental role in establishing boundaries within living systems, from the intracellular level to collectives of organisms (*1-4*). In early embryos, cell boundaries are established by rapid cleavage divisions that robustly organize the cytoplasm into progressively smaller cellular compartments (*5, 6*). Strikingly, the compartmentalization of the cytoplasm can occur before (*7*) or without (*8, 9*) the formation of a new plasma membrane, raising the question of how boundaries between cytoplasmic compartments can be robustly maintained in the absence of physical barriers. Experiments using reconstituted cytoplasm have revealed that cytoplasmic compartments self-organize spontaneously (*7, 8*). The formation and division of these compartments rely on microtubule asters that define their boundaries (*10-13*) and dynein activity that transports organelles towards the compartment center (*14*). Microtubule asters grow via self-amplifying microtubule growth or autocatalytic nucleation, the nucleation and branching of microtubules from existing ones (*12, 15*). Through branching nucleation, asters can explore a large volume of cytoplasm until they meet other asters. When asters enter in contact, the aster-aster interface is thought to be stabilized by components that provide local inhibition to microtubule nucleation and growth, creating robust boundaries that guide cytokinesis (*16-18*). This process leads to a regular tessellation of the cytoplasm similar to Turing patterns (*19*). However, it is unclear how local inhibition in combination with autocatalytic growth can lead to stable and robust boundaries (*20, 21*). To shed light on this problem, we combine theory with experiments in reconstituted cytoplasm and living embryos of zebrafish and *Drosophila*. Starting form a theoretical prediction, we show that microtubule autocatalytic nucleation gives rise to aster invasion driving the coarsening of cytoplasmic compartments. By performing cell cycle perturbations and biophysical measurements of microtubule dynamics, we find that coarsening of cytoplasmic compartments is prevented either by synchronizing the cell cycle oscillator to the dynamics of the asters or by reducing autocatalytic nucleation. Finally, we show that these mechanisms yield divergent cytoplasmic organization strategies in embryos.

## Cytoplasmic partitioning by autocatalytic microtubule waves is intrinsically unstable

We investigated cytoplasmic partitioning in live zebrafish embryos and *X. laevis* frog egg extracts. In zebrafish embryos, cytoplasmic partitioning occurs prior to cytokinesis. During the first rounds of cell division, microtubule asters divide the cytoplasm into two cytoplasmic compartments before the cell membrane ingresses (**Fig. 1A**). Moreover, the system remains syncytial until the 32 cell stage. This observation led us to test if the embryo can divide its cytoplasm in the absence of cytokinesis. To this end, we inhibited the formation of cleavage furrows by adding Cytochalasin B (*22*), an actin polymerization inhibitor. We observed low-density regions of microtubules and cytoplasmic actin between compartments over multiple cell cycles, indicating that the division of the cytoplasm in living zebrafish embryos does not require cell membranes (**Fig. 1B, Fig. S1, and video S1**). In frog extracts, undiluted cytoplasm obtained by crushing frog eggs at high speed self-organizes into distinct compartments that are not separated by cell membranes, similarly to syncytial systems (*8*). These compartments form in the absence of cytokinesis and divide over multiple cell cycles. **(Fig. 1C-D and video S2)**. These results demonstrate that cytoplasmic partitioning is a fundamental process in cell division that precedes and is independent of cytokinesis.

**Fig. 1:**
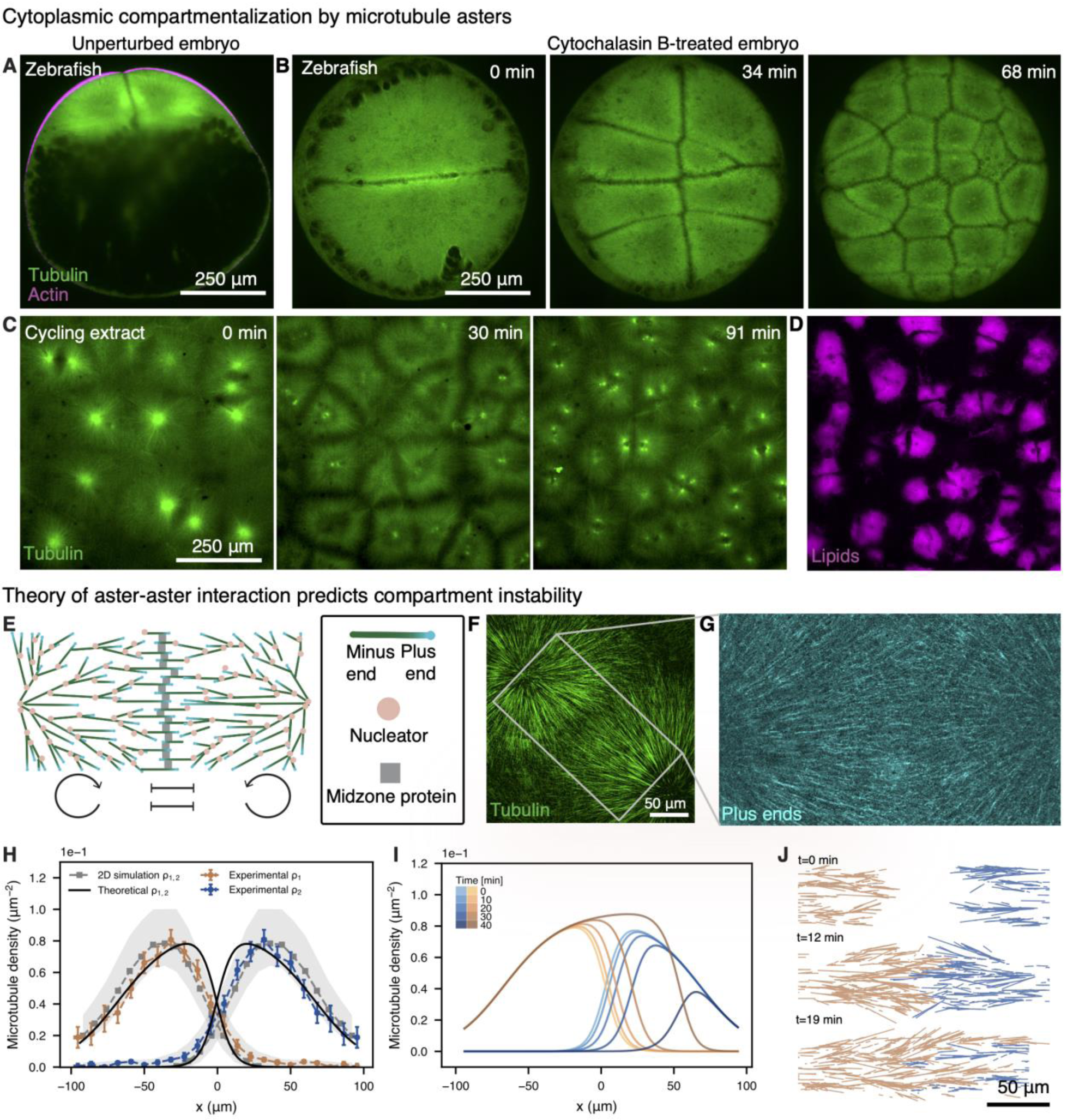
Robust compartmentalization is observed *in vitro* and *in vivo*, but theory predicts a physical instability. **(A)** Light sheet fluorescence microscopy image of a zebrafish embryo at the first cell stage. Microtubule asters form, interact and partition the cytoplasm before cleavage furrow ingression. Microtubules, labeled by EGFP-Doublecortin, are shown in green and the actin cortex, labeled by utrophin-mCherry, in magenta. **(B)** Live imaging of cytochalasin B-treated embryo where cell membrane ingression is inhibited. Asters coexist and form boundaries of low microtubule and cytoplasmic actin density. (**C**) Live imaging of cycling extract showing cytoplasmic partitioning by microtubule asters over multiple cell cycles. Microtubules are labeled by Alexa640-tubulin and shown in green. (**D**) Confocal microscopy image of cytoplasmic compartments visualized by labelling lipid organelles with Rhodamine in magenta. **(E)** Schematic of two-asters interacting. Aster-aster interaction can be described by a network of two self-amplifying loops interacting via local inhibition. (**F**) Microscopy image of two asters interacting. (**G**) Time projection of five frames of microtubule plus ends labeled with EB1-mApple, referring to the area labelled in white in (F). **(H)** Microtubule density profile of two asters as they start interacting, obtained by measuring the density of growing plus ends in the growth direction in the region where the asters interact. *x* indicates the linear coordinates from the region close the center of one aster to the region close to the center of the adjacent aster. Experimental data are shown by the points in blue and orange, agent-based simulations are reported in gray, and the one-dimensional theory is plotted in black. (**I)** Numerical time evolution of the theoretical microtubule densities shown by the black line in (H). The time evolution shows that the system is unstable as the left aster invades the right one in less than one hour. (**J)** Agent-based simulation of two interacting asters in a planar two-dimensional channel.

The striking similarities in cytoplasmic partitioning between frog egg extracts and live embryos suggest that extracts are a prime system to investigate this process, as it has the advantage that it is easy to manipulate and image. To quantify the formation of compartment boundaries we measured the microtubule density profile using EB1-mApple, as it labels the growing plus ends of microtubules (**Fig. 1 E-G**). We tracked individual EB1-mApple comets and reconstructed the density and polarity for microtubules of two adjacent compartments (**Fig. 1H and SI2**). The two profiles corresponding to each compartment have an exponential increase close to their center consistent with autocatalytic growth. Near the interface, the microtubule profiles decay consistent with local inhibition at the antiparallel microtubule overlap (*16, 17*). These profiles suggest that the interaction between the two asters can be minimally described by a network of two autocatalytic, or self-amplifying, loops interacting via local inhibition (**Fig. 1E, bottom**). To test whether such a network can explain the robust formation of these compartments, we used a 1D continuum theory of aster-aster interaction incorporating autocatalytic growth, microtubule polymerization and turnover (*15, 23*), and local inhibition (**SI**), resulting in the following equation:

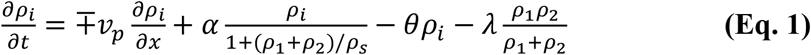

where *i =* 1,2 and refers to the two asters, *ν*_*p*_ is the polymerization velocity, *θ* the microtubule turnover, *α* is a parameter related to the autocatalytic growth, *λ* modulates the inhibition between asters, and *ρ*_*s*_ is a density of microtubules that indicates the saturation of microtubule nucleation due to depletion of nucleators as they bind to microtubules (**SI**). We measured *ν*_*p*_ by tracking the plus ends of the microtubules and *θ* using single molecule microscopy of sparsely labelled tubulin dimers. To estimate *α*, we used the initial explosive growth of microtubules close to the center of the compartments, far from the interaction zone between asters. Finally, we estimated the local inhibition *λ* by measuring the slopes of the density of microtubules at the interaction zone. A detailed description of the measurements and estimations of the parameters is reported in the SI. With all parameters fixed, we predicted the aster density profiles (**Fig. 1H, black line**). Using the measured parameters, we also validated our continuum theory using agent-based simulations **(Fig. 1H, gray lines, and Fig. S3 and S4**). Although we found quantitative agreement between the experimental and predicted profiles, both theory and simulations predict that the temporal evolution of these boundaries is unstable (**Fig. 1I-J, and video S3)**, which was not observed in the cytoplasmic extracts or embryos in **Fig. 1A-D**. Even though not observed experimentally, this instability is generally expected from local inhibition and self-amplification alone (*20*).

One possible explanation for this apparent inconsistency between theory and experiments is that the time needed to develop such instability may be larger than the cell cycle time, which drives the disassembly of the microtubule asters prior to the assembly of mitotic spindles. Close to the unstable point, the time to develop the instability can become arbitrarily large. Indeed, our numerical solutions suggest that the time to develop this instability can easily be up to 40 min (**Fig. 1I**), which is comparable to the cell cycle time in both frog extracts and frog embryos which are equal to about 40 minutes and 30 minutes, respectively (*24, 25*). To test whether the cell cycle prevents the development of the instability, we arrested cytoplasmic extracts in interphase by blocking translation of cyclin B1 with cycloheximide (*8*) **(Fig. 2A and B)**. In this condition, we observed that compartments that initially formed with a well-defined boundary as in the control condition, started coarsening by means of the microtubule asters invading each other, consistent with the aster invasion predicted by our theory. Compartment invasion was also accompanied by disassembly of the chromosomal passenger complex at the aster-aster interface (**Fig. S5 and video S4**). This coarsening continued for several hours, leading to compartments of few millimeters in size, in contrast to hundreds of microns in the cycling extract. During the coarsening, dynein motors kept relocating nuclei to the new center of the larger compartments (**Fig. 2A, bottom)**. With a closer examination using higher resolution imaging (**video S5**), we observed that an invading aster gained mass, consistent with continuous autocatalytic growth, at the expense of the invaded aster, that eventually disappeared **(Fig. 2B)**. This invasion dynamics was also reproduced in the agent-based simulations **(Fig. 2C)**. Altogether, our results show that cytoplasmic partitioning is an intrinsically unstable mechanism.

**Fig 2:**
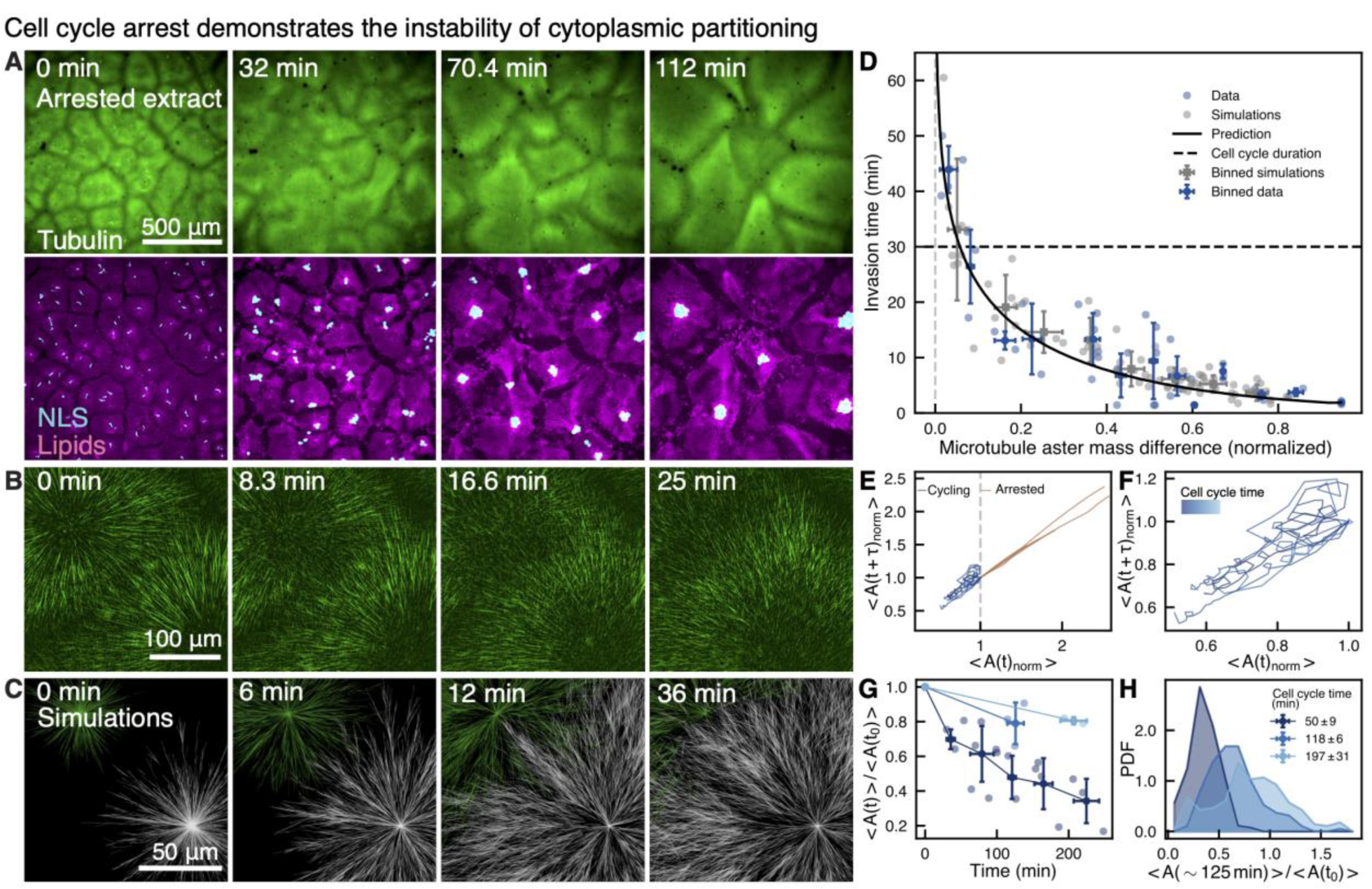
Cytoplasmic partitioning is intrinsically unstable, but the cell cycle duration can avoid the instability leading to robust compartmentalization. **(A**) Live imaging of interphase-arrested cytoplasmic extract showing microtubule aster invasion. Microtubules are shown in green. Aster invasion results in the coarsening of cytoplasmic compartments and dynein-induced relocation of sperm nuclei. Cytoplasmic compartments are shown in magenta and nuclei in cyan labelled by GFP-NLS. (**B**) High-resolution time lapse of an invasion event. The two asters compete for mass and while the invading aster gains mass nucleating and polymerizing more microtubules, the invaded aster loses mass via microtubule turnover. **(C)** Agent-based simulations of two asters showing the invasion process over time. (**D**) Invasion time plotted against initial mass difference between the asters, showing that asters with small mass differences take longer to invade than asters with large mass differences. Cell cycle time of the frog embryo (*25*) is plotted for comparison. Simulations are shown in gray, experimental data in blue, and theory in black. **(E)** Phase portrait of the average area of the compartments for arrested (orange) and cycling (blues) extract normalized for the initial area equal to 1. *τ* is equal to 8 minutes. While the area of compartments in arrested extract grows freely, the area of compartments in cycling extract oscillates and maintains a small size, even though some invasion events are present. **(F)** Zoomed graph of the phase portrait of the cycling extract. Darker colors represent shorter cell cycles. Average cell cycle time varies from 39 to 65 minutes. **(G)** Normalized average compartment area over time. Shades of blue refer to the different cell cycle times reported in the legend of (H). **(H)** Probability density function of normalized average compartment area.

## Cell cycle duration controls the size of cytoplasmic compartments and can prevent their instability

Our results suggest that the cell cycle duration can determine whether invasion events occur and thus regulate the patterns of cytoplasmic partitioning. To further investigate this dependence, we experimentally quantified the invasion time as a function of the aster mass difference, ΔM_*i*_. We calculated the mass of the asters from the area under the curve of one-dimensional profiles of microtubule density (**Fig. S6**). We defined the invasion time *τ* as the time for the initial mass difference between the asters ΔM_*i*_ to decrease by a factor *e* (**Fig. 2D)**. The invasion time decays as the mass difference between the asters increases. This trend is also perfectly captured by a parameter-free prediction of the theory (black line) and agent-based simulations (gray dots and **Fig. S7**). Asters with large mass differences invade in a few minutes. Interestingly, asters with small mass differences—that represent the situation in living embryos where compartments are highly uniform—have an invasion time comparable to the cell cycle time.

The time dependence of the invasion events allows the cell cycle duration to prevent invasion events, and therefore the runaway growth of asters, if compartments are similar in size. Conversely, for slow cell cycle times, invasion events may lead to increasing differences between compartments that may amplify the instability, leading to divergent compartment size distributions. This process can be visualized by means of a phase portrait, showing that while in the arrested extract the compartment size monotonically increases, in the cycling extract it oscillates around a characteristic compartment size, despite some invasion events (**Fig. 2E-F**). To further explore the effect of the cell cycle duration on the compartment size, we systematically delayed the cell cycle time by titrating cycloheximide amounts in extracts. These experiments showed that the cell cycle duration directly affects the average compartment size and therefore patterning of the cytoplasm (**Fig. 2G and video S6)**. Moreover, we observed that while for shorter cell cycle times the distribution of compartment sizes is narrowly peaked, as the cell cycle slows down the compartment size distribution becomes increasingly broader (**Fig. 2H)**. These results show that a delicate balance between the cell cycle time and compartment growth is necessary to achieve a uniform and robust cytoplasmic partitioning.

## The interplay between microtubule turnover and autocatalytic growth regulates the stability of compartment boundaries

Our data shows that changes in cell cycle timing can have dramatic consequences in the precision of cytoplasmic partitioning, from extremely regular partitioning when matching autocatalytic growth, to system-size coarsening. We wondered if there were regimes in the parameter space that could prevent this instability, independently of the cell cycle timing. To this end, we performed a linear stability analysis of **Eq. 1**, leading to the stability criterion:

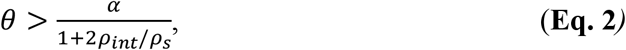

where *ρ*_*int*_ is the density of microtubules where the two asters intersect. We also confirmed this stability criterion by numerically solving **Eq. 1** (**Fig. 3A and S9**). Surprisingly, the stability of the compartment boundaries critically depends on a competition between the autocatalytic rate and microtubule turnover, and not on the strength of local inhibition (**SI**). When the autocatalytic term dominates over turnover, microtubule density profiles feature exponential growth from the center of the compartment (**Fig. 3B (ii)**). Although this density will go down as the asters interact, the boundary they form will always be unstable. Conversely, if turnover dominates over autocatalytic growth, the density of microtubules decreases from the center of the compartment (**Fig. 3B (i) and Fig. S8 for time evolution**). In this regime, the boundary created as the two asters interact will be stable, but the compartments will be generally smaller with a size defined by the decay length scale of the microtubule density. Consistent with the instability we measured, extracts fall in the unstable region of the phase diagram (**Fig. 3A)**. Although not observed in our system, these results show that the stability of compartments can be achieved by modulating microtubule nucleation and dynamics, independently of cell cycle timing.

**Fig. 3:**
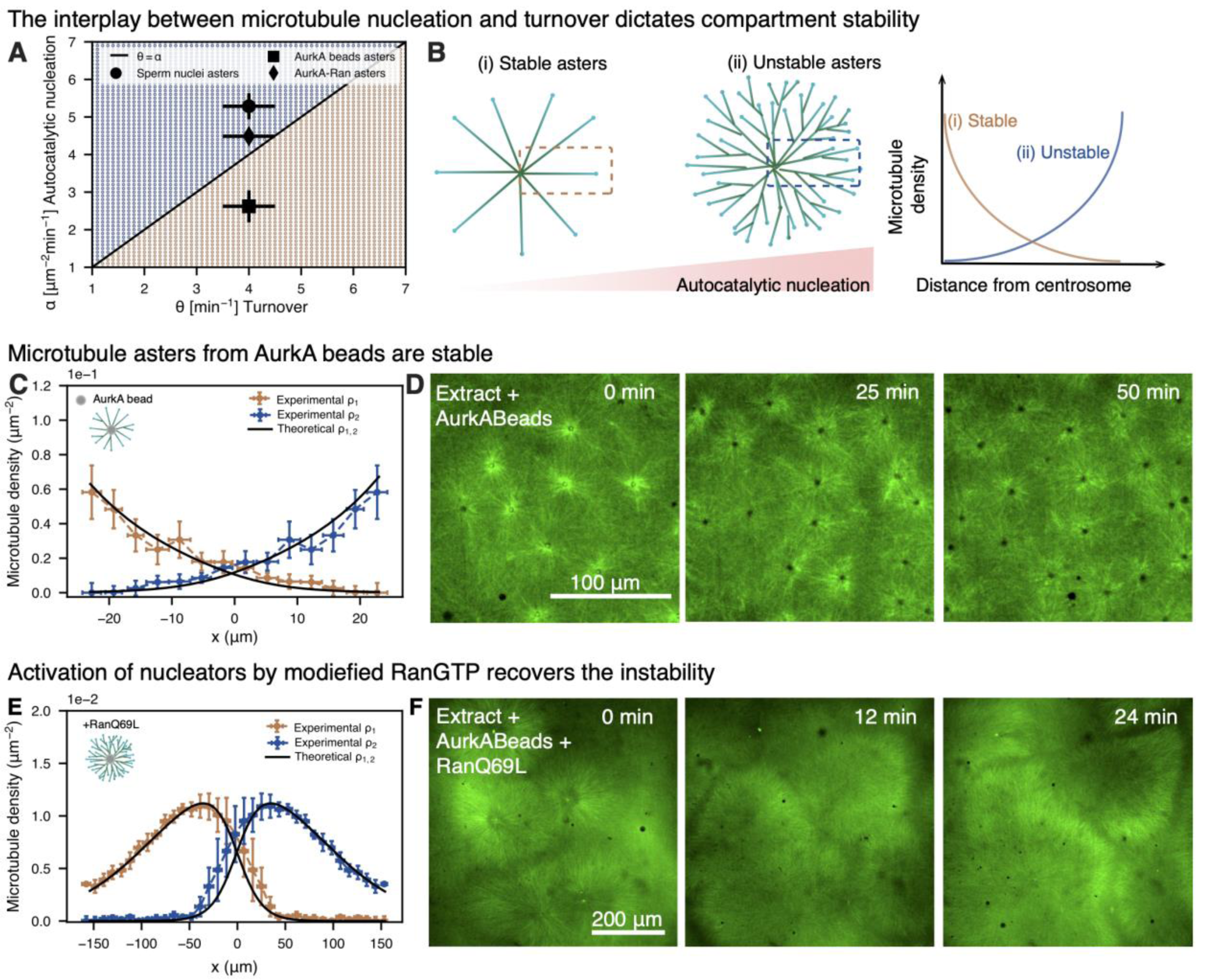
Microtubule dynamics can regulate the stability of cytoplasmic partitioning. **(A)** Phase diagram of *α* and *θ* showing a stable and unstable region. The blue and orange dots correspond to numerical solutions of **Eq. 1** with steps of 0.1 for *α* and 0.1 for *θ*. The black line represents the stability criterion. (**B**) Schematics of microtubule asters and their one-dimensional density from close to the center to the boundary depending on the amount of autocatalytic nucleation. In asters with low autocatalytic nucleation, the microtubule density decreases from the center. In asters with high autocatalytic nucleation, the microtubule density increases from the center because of the exponential nature of branching. **(C)** Microtubule density profile of two AurkA asters measured as density of plus ends of microtubules. Schematic of AurkA asters on the top left. (**D)** Confocal microscopy time sequence of AurkA asters in interphase-arrested cytoplasmic extract showing that the asters are stable and regularly partition the cytoplasm. **(E)** Microtubule density profile of two AurkA-RanQ69L asters measured as density of plus ends of microtubules. Schematic of AurkA-RanQ69L asters on the top left. (**F**) Confocal microscopy time sequence of AurkA-RanQ69L asters in interphase-arrested cytoplasmic extract showing that the asters are unstable.

To investigate the possibility of stabilizing cytoplasmic compartments by changing microtubule dynamics, we fabricated asters with a decreasing microtubule density profile (*12*). These asters can be obtained by adding Aurora kinase A-coated (AurkA) beads to extracts (**Fig. 3C**) instead of sper nuclei. The AurkA beads act as artificial centrosomes (**Fig. 3C, schematic)**. AurkA beads trigger the nucleation of microtubules (*12, 26*), but to a lesser extent than with chromatin-associated centrosomes. In this condition, we measured that the microtubule density profile decays from the beads (**Fig. 3C**), consistent with a decrease of microtubule nucleation and a stable system according to the theory. We confirmed this shift to the stable regime by measuring the nucleation and turnover rates. As expected, these values fall into the stable regime in the phase diagram (**Fig. 3A, orange area and Figure S8 for time evolution)**. We then tested if the system is stable when the cell cycle is arrested. As predicted by the stability criterion, asters formed by addition of AurkA beads in arrested cytoplasm do not invade, and end up partitioning the cytoplasm with surprising regularity similar to asters in *Drosophila* extract and embryos (*27*) (**Fig. 3D and video S7**). These results are consistent with previous experiments performed in extract with AurkA beads (*16*). To confirm that this effect was solely due to changes in microtubule nucleation and not the use of artificial centrosomes, we supplemented extracts in the presence of AurkA beads with constitutively active RanQ69L to increase microtubule nucleation (*15*) (**Fig. 3E**). In this situation, the density of microtubules from the center of the compartments increased similarly to control situation (**Fig. 3E**). Moreover, AurkA-RanQ69L asters became unstable and invaded as in the control case (**Fig. 3F and video S7**), consistent with theory. In summary, robust compartmentalization of the cytoplasm can be achieved in a parameter regime where microtubule turnover dominates over autocatalytic nucleation rate, independently of the cell cycle time.

## Regulation of microtubule nucleation captures divergent strategies of cytoplasmic partitioning in early development

To investigate the *in vivo* relevance of the stability prediction, we turned to zebrafish and *Drosophila* embryos. We chose these embryos because of their drastically distinct aster structure despite a comparable embryo size (∼700 μm in diameter for zebrafish and ∼500 μm for the long axis of *Drosophila*). In zebrafish embryos, the density of microtubules in interphase asters increases from the centrosome until microtubules reach the entire cell (**Fig. 4A and D and video S8**). In contrast, in *Drosophila* embryos, microtubule density decreases from the centrosomes (**Fig. 4B and F and video S8**) and microtubule asters do not reach the boundary of the whole syncytium (cortex of the embryo). These asters slowly fill up the embryo volume in subsequent cell divisions. Based on the theory and results in extract, we predict that the cytoplasmic compartments in zebrafish should be unstable and by contrast in *Drosophila* stable. To test this prediction, we first confirmed where these embryos lie in the phase diagram (**Fig. 4C**). To this end, we quantified microtubule dynamics in embryos by measuring the polymerization velocity as the speed of plus ends, and the microtubule turnover as half time recovery from photobleaching (for zebrafish) and photoconversion experiments (for *Drosophila*). We estimated the parameters associated to autocatalytic growth and the local inhibition similarly to the data of microtubule asters in extract. In the stability phase diagram, zebrafish falls into the unstable region whereas *Drosophila* lies in the stable region, consistent with the shape of the density profiles. Interestingly, the microtubule turnover we measured in extracts, zebrafish, and *Drosophila*, is very similar, while the shift from the stable to unstable regime is mainly driven by changes in microtubule nucleation.

**Fig. 4:**
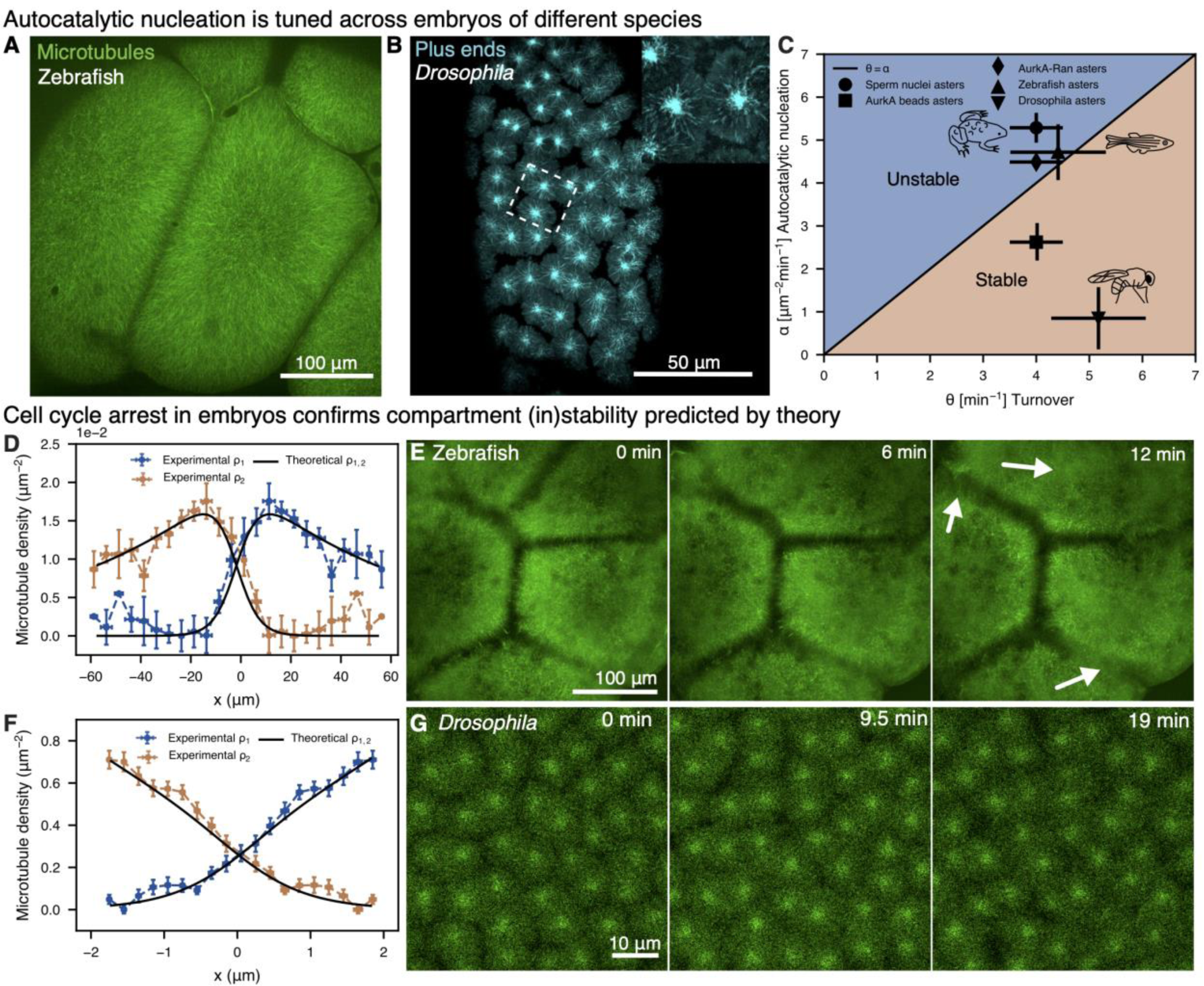
Test of the (in)stability prediction in zebrafish and *Drosophila* embryos. **(A)** Confocal microscopy image of two microtubule asters in zebrafish. **(B)** Confocal microscopy image of microtubule asters in *Drosophila* visualized by a time projection over 20 frames of growing plus ends shown by EB1 using a transgenic line. In the inset, zoomed image of the growing plus ends. **(C)** Phase diagram of *α* and *θ* including the data of zebrafish, *Drosophila*, and frog cytoplasm. **(D)** Microtubule density profile of asters of zebrafish. The microtubule density profiles increase from the center of the aster indicating that aster growth is dominated by autocatalytic nucleation. (**E)** Live confocal imaging of interphase-arrested zebrafish at cycle 3 showing invasion events with use of white arrows. **(F)** Microtubule density profile of asters of *Drosophila*. The microtubule density profiles decrease from the center of the aster indicating that aster growth is dominated by turnover. (**G)** Live imaging of interphase-arrested *Drosophila* at nuclear cycle 12 showing aster stability.

We next tested the stability prediction by arresting the cell cycle in interphase *in vivo* by adding cycloheximide, and following the aster dynamics using live imaging. We arrested the cell cycle at cycle 3 and nuclear cycle 12 for zebrafish and *Drosophila*, respectively. As predicted, zebrafish compartments were unstable and invaded each other within 15 minutes (**Fig. 4E** and **video S9**). As in extracts, the invasion events drive the dynein-mediated re-localization of nuclei (**Fig. S10**), strongly affecting the cytoplasmic organization in the embryo. In contrast, compartments in *Drosophila* remained stable, reminiscent of the compartments formed in extracts with AurkA beads (**Fig. 4G** and **video S9)**. Because AurkA beads resemble the compartmentalization of *Drosophila* embryos, we wondered if changing microtubule nucleation alone not only dictates the stability of the compartments but also the dynamics of organization of the entire cytoplasmic volume as in *Drosophila* embryos. To test the “*Drosophilization”* of the extract, we looked for regions in the cytoplasm where there were only centrosomes in the absence of DNA. The centrosome asters had similar profiles to the AurkA beads and *Drosophila* embryos. These asters progressively filled the volume as they divided, similarly to *Drosophila* embryos (**Fig. 5A** and **video S10)**, and with stark contrast to the complete covering of the whole cytoplasm in control extract during each cell cycle (**Fig. 5B** and **video S10)**. Altogether, our data show that our stability criterion can predict the dynamics of divergent compartmentalization strategies *in vivo*, which can be explained by tuning the amount of autocatalytic microtubule nucleation. In frog and zebrafish, microtubule asters grow with high autocatalytic nucleation which leads to large asters that can reach the embryo boundary and therefore cover the whole embryo cytoplasm from the first cell stage, but are unstable. In *Drosophila*, where asters possess low autocatalytic nucleation, compartments are stable, but small, and fill the cytoplasm over multiple divisions, leading to lower cytoplasmic coverage (**Fig. 5C-D)**.

**Fig. 5:**
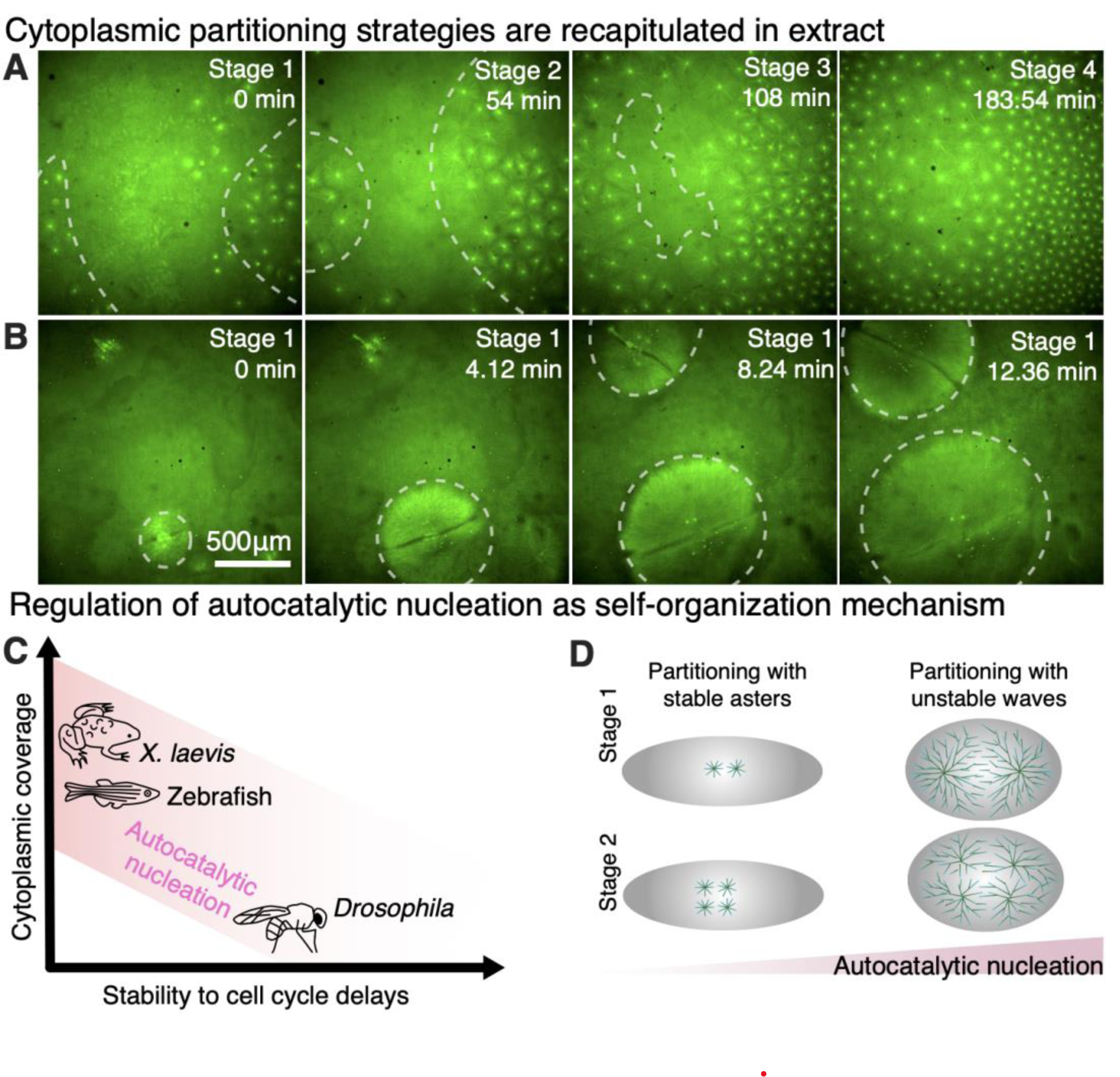
Divergent strategies of cytoplasmic partitioning driven by regulation of microtubule self-amplification. **(A**) Microtubule asters in cytoplasmic extract with centrosomes and slower cell cycle time. Centrosomal nucleation without chromatin gives rise to asters with a decaying microtubule density profile that can fill the cytoplasm over multiple cell cycles, similarly to the first stages of the *Drosophila* embryo. Slower cell cycle is chosen here to highlight the filling process, which for centrosomal asters also occurs at control cell cycle times. **(B**) Microtubule asters in cytoplasmic extract with sparse sperm nuclei where asters grow to mm-size in one cell cycle. (**C**) Phase diagram of cytoplasmic organization in early embryos. To organize the cytoplasm before cellularization and over large scales, early embryos tune the level of autocatalytic growth. Frog and zebrafish embryos start cellularizing at the first division, requiring microtubule asters to rapidly grow mass and cover the entire embryo cytoplasm. The disadvantage of this strategy is that it is unstable and it needs to be timely controlled by the cell cycle oscillator. In *Drosophila* embryos, cellularization occurs after 13 nuclear divisions, therefore asters do not require to exploit high levels of autocatalytic growth. As a result, asters in *Drosophila* embryos are small and stable and gradually cover the cytoplasm over multiple divisions. **(D)** Schematic of these divergent strategies to organize the cytoplasm.

## Discussion

Using a theory for aster-aster interactions, and experiments in extracts and *in vivo*, we revealed that cytoplasmic partitioning prior to cytokinesis is an intrinsically unstable mechanism in large vertebrate embryos. This instability originates from a competition between microtubule autocatalytic growth and turnover. Despite this inherent instability, we demonstrate that precise cell cycle timing renders this compartmentalization dynamically stable, resulting in remarkably robust partitioning of the cytoplasm. To find the proper geometric center, cells read the geometrical boundaries of the embryo using unstable microtubule waves that reach the cortex (*10, 28*). This instability imposes a delicate balance between the waves of autocatalytic growth and the cell cycle timing. The cell cycle duration needs to be slow enough for waves to read the geometry of the cell but fast enough that compartments do not fuse. The cell cycle also needs to be synchronous across compartments to avoid invasion events, as seen in zebrafish embryos and *Xenopus* egg extracts. In organisms that do not require early cellularization, as in syncytial *Drosophila* embryos, it is not necessary to immediately read the cell geometry. Instead, smaller and stable asters that compartmentalize a shared cytoplasm can slowly divide and fill up the embryo space during 13 cell cycles prior to cellularization. In this situation, there is no need to rely on unstable autocatalytic processes or to have a perfectly synchronized cell cycle as the compartments remain stable.

Our study underscores that the diverse compartmentalization behaviors observed across species can be explained by the interplay between microtubule turnover and nucleation. As turnover remains conserved among the species examined, our findings suggest that evolutionary changes in microtubule nucleation may contribute to the diverse cytoplasmic partitioning strategies across species. These findings are crucial not only in the context of embryonic development but also for syncytial systems and cytokinesis. In syncytial systems, where cytoplasmic compartments lack cell membrane separation, mechanisms regulated by the cytoskeleton are essential for maintaining distinct borders. Similarly, during cytokinesis, cells briefly become syncytial and must sustain separate cytoplasmic compartments until cytokinesis is completed (*7*).

This work presents a novel integration of Turing-like mechanisms with biological oscillators, contributing to the understanding of pattern formation dynamics. We explore a network characterized by local self-amplification (autocatalytic growth of the asters) and local inhibition. This network is unstable, however, when properly modulated with the cell cycle oscillator, it gives rise to dynamically stable and robust states. This combination of unstable networks with oscillators unlocks a realm of previously unexplored unstable regimes, yielding dynamically stable patterns endowed with remarkable traits such as rapid spatial coverage and flexibility. Departing from the traditional characterization of robust Turing networks as stable (*21, 29*), our study illuminates the potential of coupling unstable mechanisms with oscillators.

Overall, our research exemplifies how precise temporal tuning of biological oscillators can govern spatial patterning and size (*30*), highlighting how physical and geometrical constraints influence the evolution of self-organization mechanisms.

## Acknowledgments

We thank Keisuke Ishihara for the RanQ69L protein and the AurkA antibody. We thank Maria Elsner for labelling the INCENP antibody. We thank Heino Andreas (MPI-CBG, frog facility) for maintaining the frogs, the fish facility (MPI-CBG), and the light microscopy facility (MPI-CBG) for support with the microscopy imaging. We thank Julia Eichhorn for the illustrations of the model organisms. We thank for Argo Mukherjee for initial discussions. We thank Dan Needleman, Martin Loose, Bruno Vellutini, and Micheal Riedl for input on the manuscript.

## Author contributions

MR, AK, and YX conducted experimental research on *Xenopus* egg extracts, zebrafish embryos and *Drosophila* embryos, respectively. MR and JB conceived the work and wrote the manuscript. BD conducted agent-based simulations. MR analyzed experimental data. JB conducted theoretical modeling, fitted the data, and supervised the work. All authors contributed ideas and reviewed the manuscript.

